# Molecular parallelism in fast-twitch muscle proteins in echolocating mammals

**DOI:** 10.1101/244566

**Authors:** Jun-Hoe Lee, Kevin M Lewis, Timothy W Moural, Bogdan Kirilenko, Barbara Borgonovo, Gisa Prange, Manfred Koessl, Stefan Huggenberger, ChulHee Kang, Michael Hiller

## Abstract

Detecting associations between genomic changes and phenotypic differences is fundamental to understanding how phenotypes evolved. By systematically screening for parallel amino acid substitutions, we detected known as well as novel cases (Strc, Tecta, Cabp2) of parallelism between echolocating bats and toothed whales in proteins that could contribute to high frequency hearing adaptations. Interestingly, our screen also showed that echolocating mammals exhibit an unusually high number of parallel substitutions in fast-twitch muscle fiber proteins. Both bats and dolphins produce an extremely rapid call rate when homing in on their prey, which was shown in bats to be powered by specialized superfast muscles. We show that these genes with parallel substitutions (*Casq1, Atp2a1, Myh2, Myl1*) are expressed in the superfast sound-producing muscle of bats. Furthermore, we found that the calcium storage protein calsequestrin 1 of bats and dolphins functionally converged in its ability to form calcium-sequestering polymers at lower calcium concentrations, which may contribute to rapid calcium transients required for superfast muscle physiology. The proteins that our genomic screen detected could be involved in the convergent evolution of vocalization in echolocating mammals by potentially contributing to both rapid Ca^2+^ transients and increased shortening velocities in superfast muscles.

**Abbreviations:**

SR: sarcoplasmic reticulum

## Introduction

An important aspect to understanding how nature’s phenotypic diversity evolved is to detect the genomic differences that are associated with phenotypic differences [1]. Despite numerous sequenced genomes, detecting such associations remains a challenge. Convergent evolution, which refers to the repeated evolution of similar phenotypes in independent lineages, offers a paradigm to computationally screen genomes for molecular changes that evolved in parallel in these lineages and thus, could be involved in the phenotypic difference [2–5].

A fascinating example of phenotypic convergence is the ability of bats and toothed whales to use biosonar (echolocation) for navigating and hunting. Both independent mammalian lineages have evolved a highly sophisticated echolocation system that involves producing biosonar signals (calls or clicks), listening for the returning echoes, and converting this information into an auditory representation of their environment. Both lineages have converged on the ability to hear high frequency sounds, which are particularly suited for the detection of small prey. Furthermore, despite having evolved two different vocalization systems (laryngeal in bats, nasal in toothed whales), both lineages have converged on adopting a similar strategy for foraging. Bats and toothed whales are able to precisely control the rate of echolocation calls and drastically increase this rate to 160 or more calls per second in the final moments of capturing prey, a feature known as the “terminal buzz”. This allows the echolocators to receive near instantaneous feedback on the position of their prey and home in with high precision [6–8].

In order to detect genomic changes involved in the convergent hearing properties of these echolocating lineages, several studies have searched and identified parallel amino acid substitutions in candidate genes with a known hearing-related function such as Dfnb59, Otof, and prestin [9–12]. Subsequently, *in vitro* assays demonstrated that parallel substitutions in prestin, a protein known to be critical for high-frequency hearing [13], account for the functional convergence in key parameters of prestin [14]. This shows that searching for molecular parallelism in candidate genes provided insights into the convergent adaptations to high frequency hearing; however, the mechanisms underlying the convergent vocalization aspect of echolocation remain unknown.

Here, we systematically searched for parallel molecular evolution to possibly find new associations to convergent phenotypic adaptations. To this end, we systematically screened one-to-one orthologous proteins for parallel amino acid substitutions in independent lineages. This screen uncovered known as well as novel cases of parallelism between echolocating bats and toothed whales in proteins that could contribute to high-frequency hearing adaptations. Unexpectedly, we found an unusually high number of parallel substitutions between these echolocators in proteins crucial for fast muscle activity. We found that these proteins are expressed in superfast muscles of an echolocating bat and show functional convergence of the calcium storage protein Casq1. Our study raises the possibility that these proteins are involved in the evolution of superfast muscles that power the terminal buzz, and provides a general framework for using computational approaches to potentially discover the genomic basis of novel phenotypes

## Results

To systematically search for parallel molecular evolution, we applied a computational method that identifies parallel amino acid substitutions occurring in pairs of independent lineages (Figure S1). First, we computed a multiple protein alignment using the phylogeny-aware Prank method [15]. Second, we inferred the most likely ancestral protein sequence at every internal node in the phylogenetic tree using both Maximum Likelihood and Bayesian approaches. Third, considering each protein position and all possible pairs of branches, we extracted those pairs in which one or more parallel amino acid substitutions have occurred (Figures 1A, S2). The result is a list of lineage pairs that share at least one parallel substitution.

**Figure 1:**
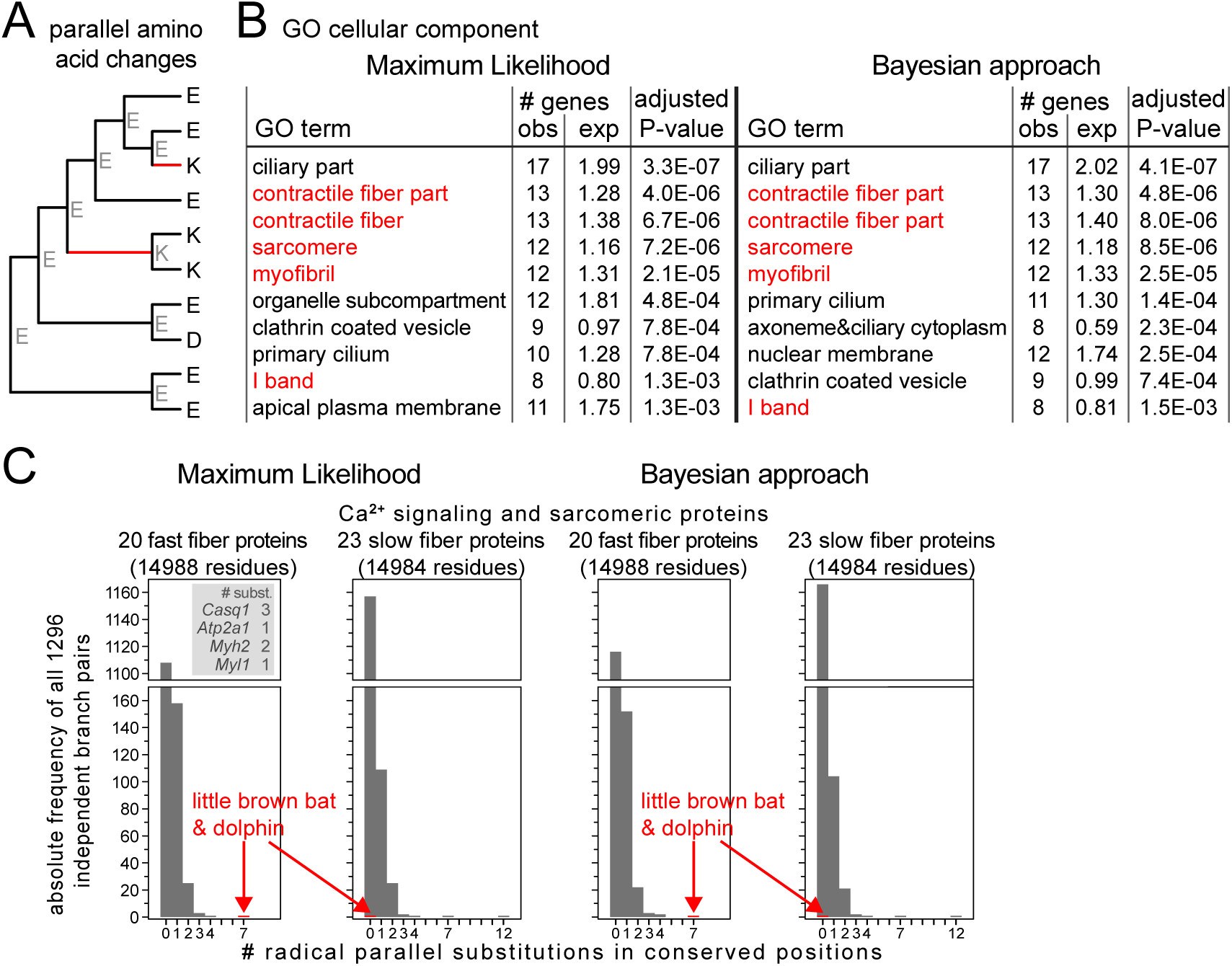
Screen for parallel molecular evolution in mammals. (A) Illustration of parallel amino acid substitutions, defined as the same amino acid substitution from an identical ancestral state. Inferred ancestral amino acids are in grey font. Red branches have a parallel substitution from E (Glu) to K (Lys) at this alignment position. (B) Functional enrichments of 409 (416) proteins, where Maximum Likelihood (Bayesian) ancestral reconstruction infers parallel substitutions between the microbat (little brown bat) and the bottlenose dolphin, using GeneTrail2 [79]. Ontology terms relating to muscle fiber components are in red font. Only the top 10 enrichments are shown for Cellular Component (65 significant enrichments). (C) Histograms showing the number of radical parallel amino acid substitutions in conserved positions of fast-twitch vs. slow-twitch Ca^2+^ signaling and sarcomeric proteins for all 1,296 independent pairs of branches. Ancestral sequence reconstruction with Maximum Likelihood (left) and a Bayesian approach (right) consistently shows that microbat and dolphin have seven parallel substitutions exclusively in fast-twitch fiber proteins (inset) and that no other branch pair has an excess of seven or more substitutions in the fast-twitch fiber set.

To systematically implement this approach, it is essential to have high quality sets of one-to-one orthologous genes that also excludes any similar paralogs. Therefore, we used the ortholog data in Ensembl Compara [16] to obtain 14,406 ortholog sets comprising of 30 placental mammals (Figure S3). Consistent with previous results [17], our screen showed that the vast majority (∼90%) of the detected parallel substitutions occur at alignment positions that are poorly conserved in evolution or are conservative replacements (similar physicochemical properties of amino acids) (Table S1). In contrast to conservative substitutions, radical substitutions that change physicochemical properties of amino acids such as charge, polarity or aromaticity occur at a lower rate in evolution, since these substitutions are more likely to affect protein function and are thus under stronger purifying selection [18, 19]. Aiming at detecting parallel substitutions that may affect function, we decided to focus on proteins with at least one parallel substitution that occurs in a conserved alignment position and that results in a radical amino acid replacement (Figure S2, Table S2).

As our dataset contained two echolocating mammals, the microbat *Myotis lucifugus* (little brown bat) and the bottlenose dolphin (*Tursiops truncatus*), we tested whether our screen detected known parallel substitutions. To this end, we extracted all proteins that contain radical parallel substitutions in conserved positions between microbat and dolphin. With Maximum Likelihood and Bayesian ancestral reconstruction, this resulted in sets of 409 and 416 proteins, respectively. For 392 proteins (>94% of 409/416), both approaches consistently detected parallel substitutions. These 392 proteins included the hearing-related proteins prestin, Dfnb59 and Slitrk6 (Figure S4A) that have been identified in previous candidate gene studies [4, 9–11].

Interestingly, our screen also detected previously unknown parallel substitutions in three additional hearing-related proteins: stereocilin (Strc), tectorin alpha (Tecta) and calcium binding protein 2 (Cabp2) (Figure S4B). The function of Strc is to connect stereocilia to each other as well as to the tectorial membrane [20], which contains Tecta as a major component [21]. Cabp2 plays a role in regulating the activity of voltage-gated calcium channels that open in response to stereocilia deflection [22]. Thus, our screen discovered additional hearing-related proteins that may contribute to convergent adaptations for high frequency hearing in echolocating mammals.

To explore the functions of the remaining non-hearing related proteins with parallel substitutions between the microbat and bottlenose dolphin, we searched for functional enrichments in these sets of 409/416 proteins. Surprisingly, we found that the proteins with parallel substitutions between microbat and dolphin are enriched for gene ontology terms relating to components of the contractile machinery of muscle fibers and calcium signaling (Figure 1B, Tables S3, S4). We examined the proteins across the different gene ontology terms for muscle and calcium signaling, and observed that while most of them also have functional roles outside of muscle fibers or are part of cardiac muscles, the following four proteins have specific roles in skeletal muscle fibers: [23, 24]: calsequestrin 1 (Casq1), ATPase sarcoplasmic/endoplasmic reticulum calcium transporting 1 (Atp2a1, synonym Serca1), myosin heavy chain 2 (Myh2) and myosin light chain (Myl1) (Figures 2 and 3). Casq1 adsorbs large amounts of Ca^2+^ ions and serves as the Ca^2+^ ion storage protein in the sarcoplasmic reticulum (SR) [25–27]. Atp2a1 pumps Ca^2+^ into the SR to achieve muscle relaxation [28]. Both Myh2 and Myl1 are components of the force-generating molecular motor complex [23, 29]. In these four proteins, we observed a total of seven radical parallel substitutions in conserved positions between microbat and dolphin.

**Figure 2:**
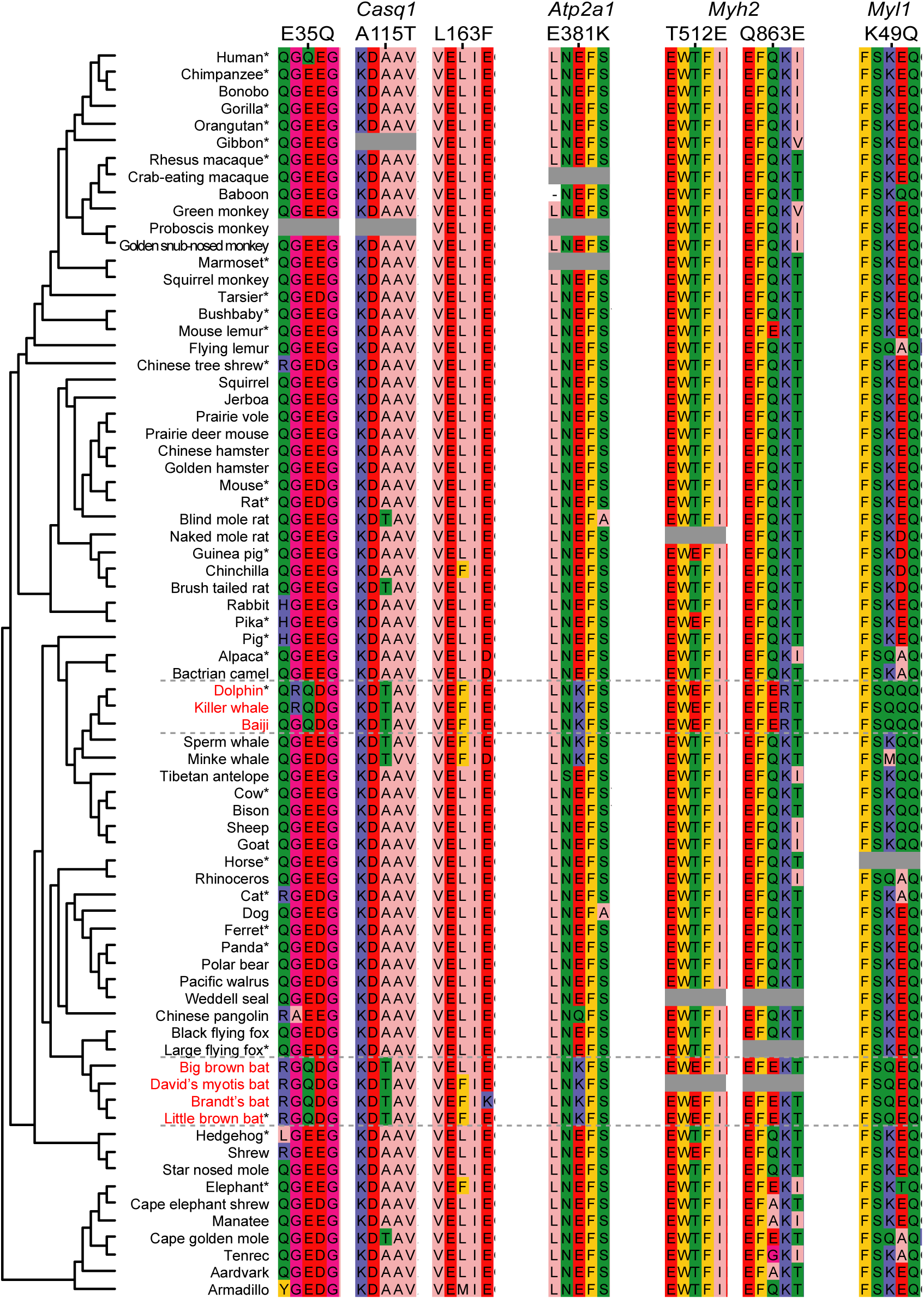
Parallel molecular evolution in fast-twitch muscle fiber proteins in echolocating mammals. In addition to species included in the genome-wide screen (marked by asterisk), the alignments contain 46 additional mammals, which shows that parallel amino acid substitutions also occur in other echolocating mammals that emit calls at a high repetition rate (big brown bat, other *Myotis* species, killer whale, and baiji) but not exclusively in them. Grey box: missing sequence.

**Figure 3:**
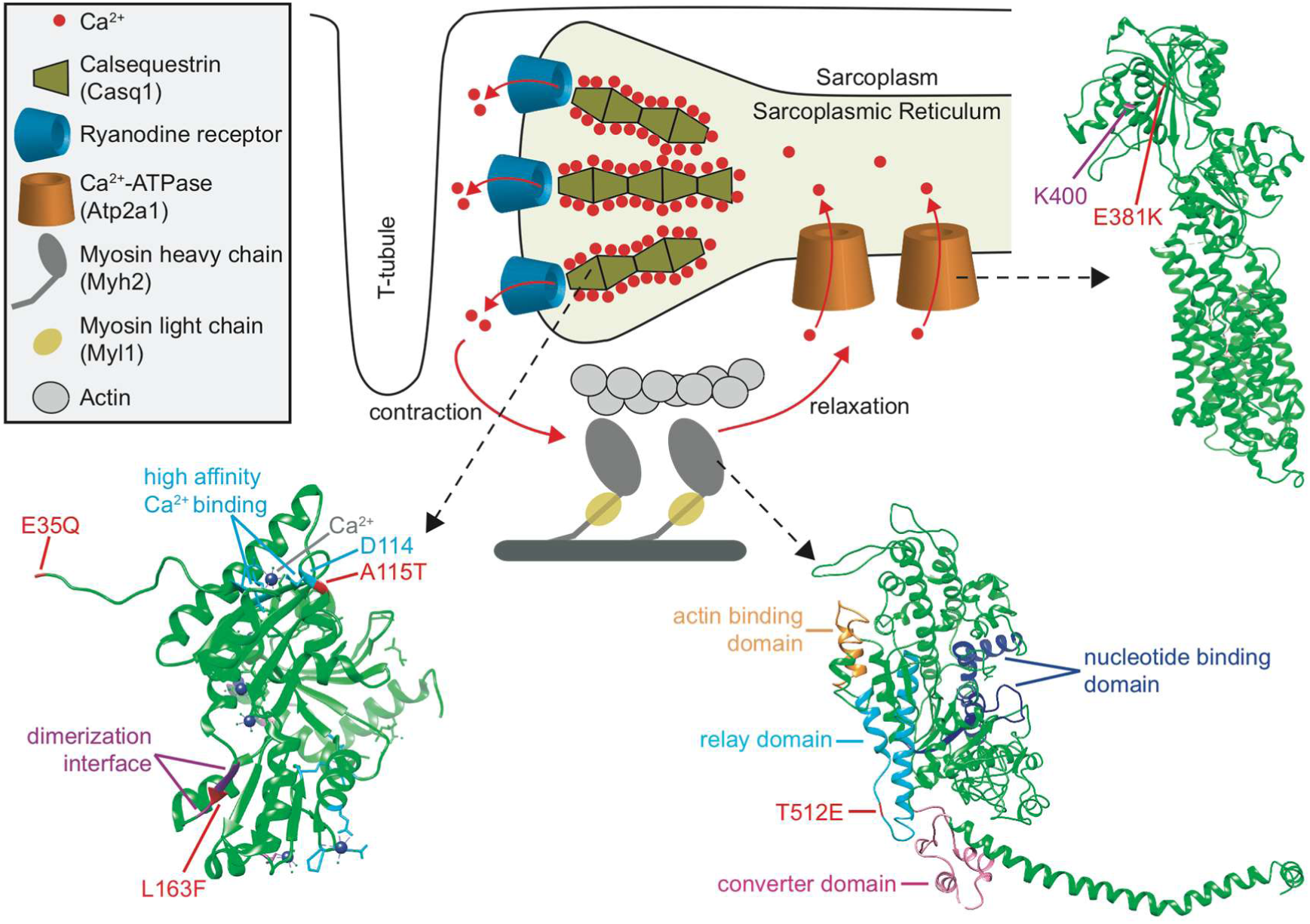
Simplified diagram showing the skeletal muscle contraction-relaxation cycle and 3D structures of proteins with parallel substitutions. Ca^2+^ bound to Casq1 is released from the sarcoplasmic reticulum through ryanodine receptors into the sarcoplasm to initiate the cross-bridge cycle during which myosin complexes generate mechanical work. For relaxation, Ca^2+^ is actively pumped back into the sarcoplasmic reticulum by Atp2a1, where majority of Ca^2+^ ions are bound again to polymerized Casq1. The 3D protein structures of Casq1 (PDB: 3TRP; residues that bind Ca^2+^ with high-affinity are in light blue), Atp2a1 (4NAB) and Myh2 (2MYS; Q863E is in the myosin light chain binding region but not part of this structure) are shown. Parallel substitutions are indicated in red. Residues involved in Casq1 dimerization and Ca^2+^ binding [27] and the Atp2a1 residue K400 that binds the inhibitor phospholamban [80] are labeled. The Myh2 relay domain that determines the actin sliding velocity [50] by communicating conformational changes between the converter domain, nucleotide-binding site, and actin-binding site [49, 81, 82] is indicated. K49Q is not part of any available Myl1 structure.

All four proteins (Casq1, Atp2a1, Myh2 and Myl1) are known to be predominantly expressed in fast-twitch skeletal muscle fibers, while respective paralogs of these proteins are usually expressed at higher levels in slow-twitch or cardiac muscles fibers. Therefore, we examined whether parallel substitutions are specific to fast-twitch muscle proteins, or whether microbat and dolphin also exhibit parallel substitutions in Ca^2+^ signaling or sarcomeric proteins that are predominantly expressed in slow-twitch muscle fibers. To this end, we used available mouse gene expression data [23, 24] and obtained 20 genes (ATP2A1, CALM3, CASQ1, MYBPH, MYH1, MYH13, MYH2, MYH8, MYL1, MYL7, MYOM2, MYOZ1, PAK1, PDLIM3, PPP3CA, PVALB, TMOD1, TNNC2, TNNT3, VCL) with a higher expression level in fast-twitch muscle fibers and 23 genes (ACTA1, ACTN2, ATP2A2, CASQ2, CRYAB, HSPB1, ITGB1BP2, ITPR1, MURC, MYH3, MYH6, MYH7, MYL2, MYL3, MYOT, MYOZ2, SMPX, SMTNL1, TNNC1, TNNI1, TNNT1, TNNT2, TPM2) with a higher expression in slow-twitch muscle fibers. In contrast to the fast-twitch muscle proteins, which included the aforementioned four proteins, we did not detect a single parallel substitution between microbat and dolphin in any of the 23 slow-twitch fiber proteins (Figure 1C). Next, we utilized our screen that detected parallel substitutions between any pair of lineages to test whether other mammals also have 7 or more parallel substitutions in the 20 fast-twitch fiber proteins. Among all other 1,295 pairs of independent branches, we observed a maximum of four parallel substitutions (Table S5), showing that the seven parallel substitutions between microbat and dolphin is an outlier observation (Figure 1C). Furthermore, while microbat and dolphin have no parallel substitutions in the 23 slow-twitch proteins, other pairs of lineages have up to 12 such substitutions in this set. This shows that the number of parallel substitutions that occur specifically in fast-twitch fiber proteins in echolocators is the highest compared to other lineage pairs. Next, we determined whether GC-biased gene conversion (a process that biases G/C over A/T alleles during recombination repair [30]) may be a potential explanation for the observed excess of parallel substitution between dolphin and microbat in the fast-twitch fiber proteins. We found that only one of the seven substitutions (K49Q in Myl1) could potentially be explained by GC-biased gene conversion (Table S6). In summary, we observed that (i) the echolocating microbat and dolphin have more parallel substitutions in fast-twitch fiber proteins than all other lineages, (ii) both species have no parallel substitutions in slow-twitch fiber proteins and (iii) GC-biased gene conversion is not a major contributing factor explaining the presence of these parallel substitutions in fast-twitch fiber proteins. While parallel substitutions often occur by chance [31, 32], these observations suggest that the seven parallel substitutions in the fast-twitch muscle proteins of echolocators, as a whole, may not have arisen due to chance alone.

In bats, the extremely rapid echolocation calls during the terminal buzz are powered by unique, sound-producing “superfast” muscles, which consist of specialized fast-twitch fibers that display contraction rates similar to the call rates during the terminal buzz (∼200 times per second) [33, 34]. Just like bats, toothed whales such as bottlenose dolphin, harbour porpoise and false killer whale use terminal buzz in the final moments before capturing prey and are able to dynamically control the echolocation click rates based on the distance to their prey [8, 35, 36]. This suggests that superfast sound-producing muscles may also exist in these toothed whales. Thus, we hypothesized that these proteins could be involved in functional changes that convert fast muscle fibers into superfast fibers, and thus potentially contribute to the convergent vocalization aspect of echolocation.

Since our orthologous gene sets comprised only 30 placental mammals including 2 echolocators, we manually analyzed 46 additional mammals to determine whether the parallel substitutions in these proteins also occurred in related echolocating bats and toothed whales. We found that most of the parallel amino acid changes are also observed in other echolocating mammals (big brown bat, additional *Myotis* species, killer whale, and baiji) though not exclusively in them (Figure 2). A further manual examination of the six aforementioned hearing-related proteins (prestin, Dfnb59, Slitrk6, Strc, Tecta, Cabp2) also shows that none of the parallel substitutions occur specifically in these echolocating mammals, with the exception of Strc H320Q that our screen uncovered (Figure S4). Notably, the N7T and P26L substitutions that were experimentally shown to affect prestin function also occur in the non-echolocating elephant shrew and Felidae (big cats) / minke whale, respectively (Figure S4A) [37]. This suggests that the observation of a derived substitution in a background species does not necessarily preclude a functional effect on the protein, and that pure patterns of convergence rarely exists when taxonomic sampling depth increases [38]. Furthermore, in various examples of such as prestin, RNAse1, rhodopsin, Na+/K+-ATPase, functional convergence is observed along with the presence of multiple sites displaying parallel substitutions [14, 39–41].

Next, we examined whether the four fast-twitch fiber proteins (Casq1, Atp2a1, Myh2, and Myl1) are expressed in the specific sound-producing superfast muscles. Although our attempts to extract RNA from the nasal muscles of the harbour porpoise (a close relative of the dolphin) were unsuccessful due to poor RNA preservation in the available samples, we succeeded in extracting high-quality RNA from the laryngeal anterior cricothyroid muscle (n=3) of the echolocating bat *Pteronotus parnellii.* Using RNA-seq, we found that all four proteins were expressed in this superfast sound-producing muscle [33] (Figure 4). Moreover, using breast muscle as control (n=3), we found that the anterior cricothyroid muscle has a substantially higher expression ratio of the four proteins relative to their slow-twitch fiber paralogs (Figure 4). We further compared expression of the three fast-twitch fiber myosin heavy chains (Myh1/2/4) that differ in their fiber shortening velocity, which decreases in the order: Myh4 > Myh1 > Myh2 [29]. We found that *Myh4* makes up 90% of the myosin heavy chain expression in the anterior cricothyroid muscle, consistent with the need for fast contraction and consistent with a recent study [42]. In contrast, Myh4 has the lowest expression in breast muscle, where the expression order of Myh4 > Myh2 > Myh1 observed in the anterior cricothyroid muscle is reversed. Interestingly, *Myh2* is the second most abundant myosin heavy chain (9%) in anterior cricothyroid muscle despite having a lower shortening velocity than Myh1. Thus, the superfast anterior cricothyroid muscle contains a higher proportion of fast-twitch fiber muscle components and expresses all four proteins with parallel substitutions.

**Figure 4:**
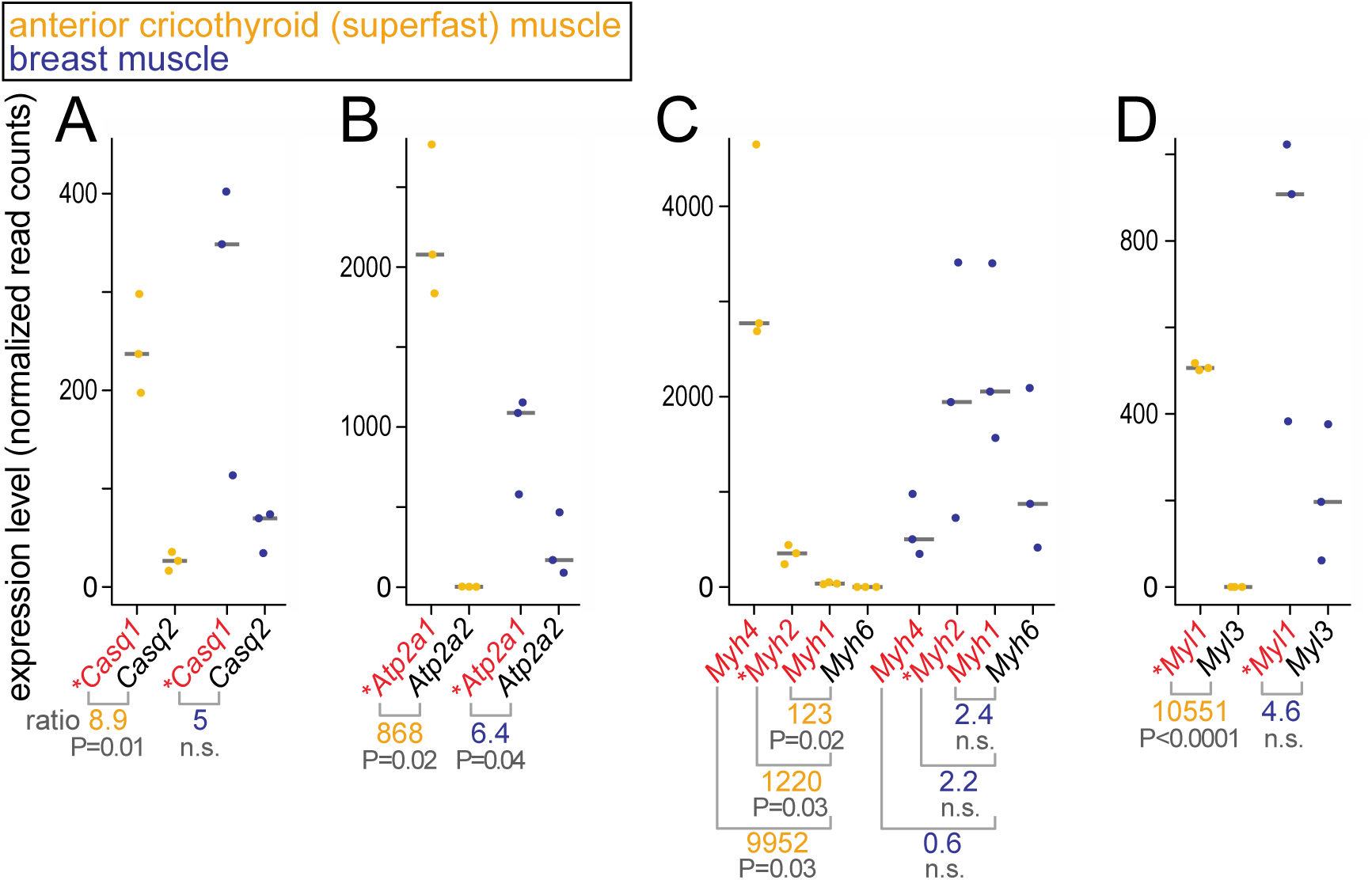
Expression level of calsequestrin (A), Ca^2+^ ATPase (B), myosin heavy chain (C) and myosin light chain (D) genes comparing fast-twitch (red font) and slow-twitch (black font) muscle fiber components in three biological replicates of *P. parnellii* anterior cricothyroid muscle (yellow) and breast muscle (blue). Horizontal line is the median. Genes with parallel substitutions are marked by an asterisk. Expression ratios and P-values of a two-sided t-test are shown at the bottom and in Table S8. n.s., not significant.

If the proteins exhibiting parallel substitutions contribute to the function of superfast fibers, we would expect that protein function has converged between microbat and dolphin in a manner that helps to achieve the extremely rapid kinetics of superfast muscles. The rate-limiting step is muscle relaxation, which involves Ca^2+^ transport from the sarcoplasm into the SR [34, 43], where it is adsorbed to Casq1. The adsorption of Ca^2+^ depends on the ability of Casq1 to form dimers and tetramers in a Ca^2+^-dependent manner, resulting in polymers that adsorb large amounts of Ca^2+^ on the negatively charged surface and between the interfaces [26, 44]. Since the superfast contraction/relaxation cycles are too short to pump all of the released Ca^2+^ back into the SR, the Ca^2+^ concentration in the SR decreases despite less Ca^2+^ being released into the sarcoplasm in subsequent cycles [45]. Therefore, Ca^2+^-dependent polymerization of Casq1 in microbat and dolphin may be adapted to the lower concentrations in the SR of repeatedly-stimulated superfast muscles.

To explore whether the parallel substitutions in Casq1 could potentially affect Ca^2+^-dependent polymerization, we examined their positions in the three-dimensional structure (Figure 3). We found that the E35Q substitution is located at the N-terminus, which participates in an arm exchange to stabilize dimer formation [44, 46]. The L163F substitution changes the hydrophobicity of the dimer interface, likely influencing the calcium-induced hydrophobic interactions. The third substitution (A115T) is located adjacent to one of the high-affinity Ca^2+^ binding residues (D114, Figure 3), which interacts with the tetrameric partner. This suggests that all three radical substitutions could affect Ca^2+^-dependent polymerization, which is crucial for the high capacity Ca^2+^ storage function of Casq1.

To experimentally investigate whether microbat and dolphin Casq1 polymerization are adapted to lower Ca^2+^ concentrations, we first conducted turbidity assays to compare Ca^2+^ concentration-dependent polymerization of Casq1 from mouse, microbat and dolphin (Figure 5A). For both microbat and dolphin Casq1, turbidity of the solution started to increase significantly at a Ca^2+^ concentration between 1.5 mM and 2.0 mM, respectively, and plateaued at ∼3 mM Ca^2+^. This is in contrast to Casq1 from mouse where substantially higher Ca^2+^ concentrations are required to increase turbidity significantly. Second, to directly monitor Ca^2+^-dependent dimerization at low Ca^2+^ concentrations, we used a multiangle light scattering experiment, which confirmed that microbat and dolphin Casq1 dimerizes at lower Ca^2+^ concentrations compared to mouse Casq1 (Figure 5B). In summary, these results demonstrate that microbat and dolphin Casq1 are able to form polymers at lower Ca^2+^ concentrations than mouse Casq1, which could facilitate Ca^2+^ adsorption under decreasing concentrations in the SR during superfast contraction-relaxation cycles.

**Figure 5:**
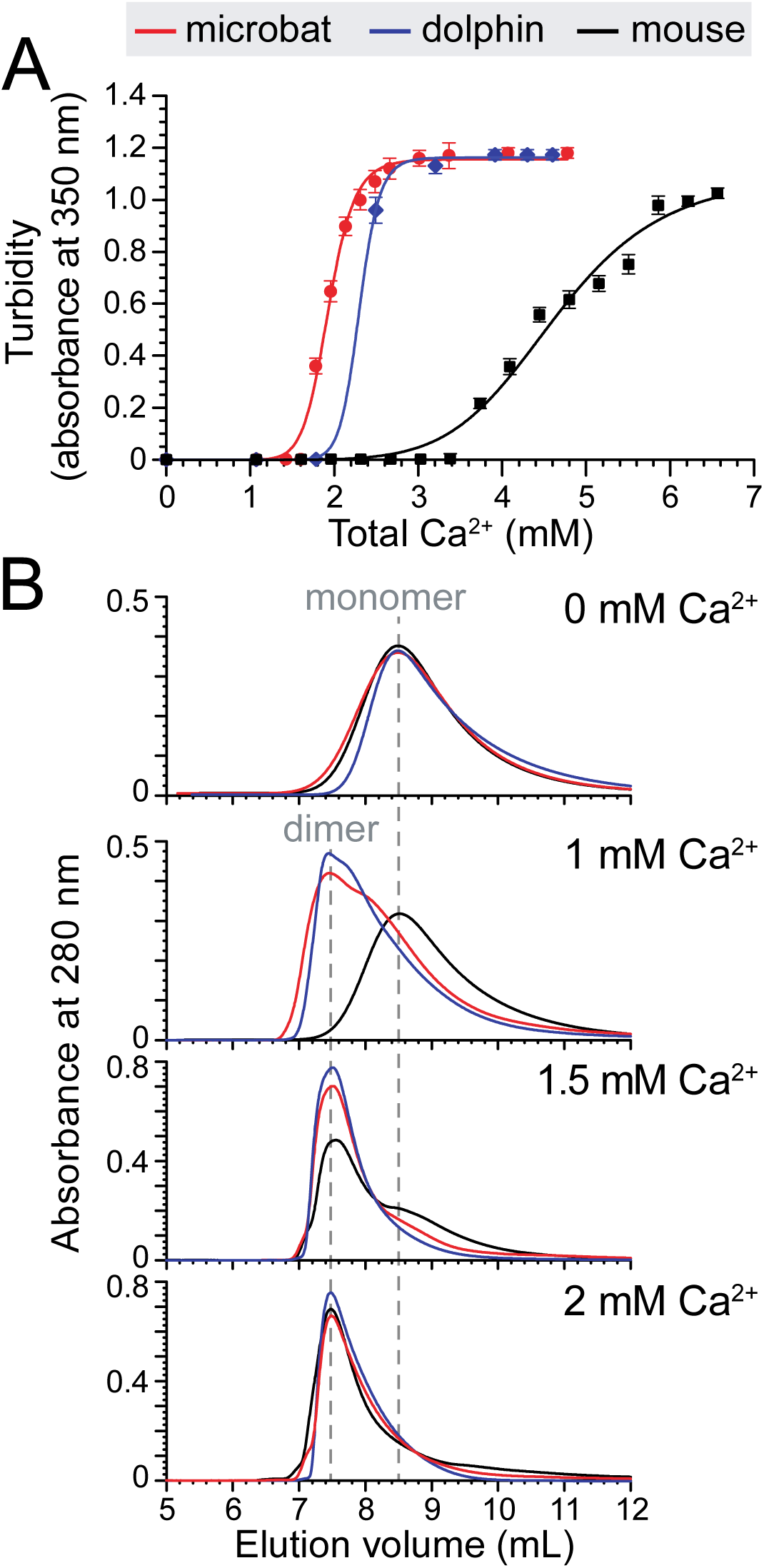
Microbat and dolphin Casq1 functionally converged in the ability to form Ca^2+^–sequestering polymers at lower Ca^2+^ concentrations. (A) Ca^2+^-dependent turbidity shows that microbat and dolphin Casq1 oligomerizes at lower Ca^2+^ concentrations. Error bars represent standard deviations of triplicate measurements. (B) Multiangle light scattering experiments show that microbat and dolphin Casq1 dimerizes at lower Ca^2+^ concentrations compared to mouse Casq1. While Casq1 from all three species is monomeric at a Ca^2+^ concentration of 0 mM, both microbat and dolphin Casq1 dimerizes at a concentration of 1 mM, in contrast to mouse Casq1 that remains monomeric.

## Discussion

Here, we show that echolocating bats and dolphins have an unusually high number of parallel substitutions in proteins that (i) are specific to the contractile machinery of fast-twitch muscle fibers and (ii) are expressed in the superfast muscle, which powers the terminal buzz in bats. Superfast muscles consist of specialized fibers capable of contracting and relaxing at a rate that is an order of magnitude higher compared to the fastest locomotor muscles [43]. To achieve this extraordinarily high rate, superfast muscles have evolved a number of key adaptations to greatly accelerate muscle relaxation, which is the rate-limiting step. These adaptations include (i) rapid Ca^2+^ transients and (ii) a higher fiber shortening velocity due to myosin motors with a fast cross-bridge detachment rate [43, 47, 48], suggesting evolutionary tinkering with various components of the Ca^2+^ signaling and molecular motor machinery. The proteins that our genomic screen detected could thus be potentially involved in the evolution of superfast muscles by contributing to both rapid Ca^2+^ transients and high shortening velocities.

Rapid Ca^2+^ transients require a fast decrease of the Ca^2+^ concentration in the sarcoplasm, which is achieved through a high amount of Atp2a1 and parvalbumin, a protein that temporarily binds Ca^2+^ in the sarcoplasm until it can be pumped back into the SR by Atp2a1 [43]. As expected, we observed a high expression level of Atp2a1 and parvalbumin in the bat anterior cricothyroid muscle (Figure 4, Table S7). In addition, our experiments show that both microbat and dolphin Casq1 are able to form polymers at lower Ca^2+^ concentrations than mouse Casq1. This functional convergence suggests that microbat and dolphin Casq1 could adsorb Ca^2+^ under conditions of lower Ca^2+^ concentrations in the SR. Such conditions are likely present in superfast muscles, where the contraction/relaxation cycles are too short to fully restore the basal Ca^2+^ concentration in the SR. Therefore, the ability of Casq1 to adsorb Ca^2+^ under lower concentrations may contribute to the rapid calcium transients required for superfast muscle physiology.

Besides rapid Ca^2+^ transients, a high fiber shortening velocity due to myosin motors with a fast cross-bridge detachment rate is also a distinguishing feature of superfast muscles [43, 48]. Fiber shortening velocity is determined by the myosin relay domain [49, 50]. Among the parallel substitutions in Myh2, the T512E substitution is located in the relay domain (Figure 3) and appears as a promising candidate to increase actin sliding velocity for two reasons. First, we found that the myosin isoform present in the Drosophila indirect flight muscles exhibits a substitution similar to T512E. These asynchronous flight muscles are capable of contracting ∼200 times per second; however, each contraction is controlled by a stretch-activated mechanism instead of a Ca^2+^ release/sequestration cycle. The myosin isoform that is exclusively expressed in flight muscles has an extremely fast cross-bridge detachment rate [48] and, as shown in Figure S5, exhibits a negatively-charged residue (Asp) at the position corresponding to Myh2 512. In contrast to the polar Thr (T), a negatively-charged residue (Glu (E) in Myh2 or Asp in Drosophila Myh) allows for the formation of a salt bridge with a positively-charged Arg in the converter domain and this interaction specifically occurs during the post-power stroke conformation that precedes cross-bridge detachment [49]. Second, mammalian Myh4 myosin, which confers the highest muscle fiber shortening velocity [29], also exhibits a negatively-charged residue (E) at position 512 (Figure S6). Therefore, the presence of a negatively-charged residue in the fastest myosins in both Drosophila and mammals supports the hypothesis that T512E is involved in increasing actin sliding velocity of microbat and dolphin Myh2.

Furthermore, we observed that the derived amino acids of additional parallel substitutions in Myh2 are also found at the corresponding Myh4 positions (Figure S6). Thus, it appears that Myh2 in microbat and dolphin have converged towards a higher similarity with Myh4. In addition, genome alignments and analysis of sequencing reads show that cetaceans have lost *Myh4* (Figures S7, S8), raising the possibility that sequence convergence in the dolphin *Myh2* compensates for the loss of *Myh4*, the gene encoding the fastest muscle myosin heavy chain.

In summary, our study highlights the utility of comparative genomic approaches to generate new hypotheses of the genomic basis underlying complex phenotypes; in this case the genomic changes that contribute to superfast muscle physiology, which has remained largely unknown. The parallel amino acid changes between echolocating mammals in several proteins, which are expressed in the fast-twitch fibers of the bat larynx, may contribute towards the functional changes required to generate exceptional speed. Even though the complete morphology of sound production in toothed whales has yet to be fully understood, the molecular parallelisms hint towards a similar role of those proteins in muscles or tissues that control sound production in these aquatic mammals. Nonetheless, additional experiments, both *in vitro* and *in vivo*, will be necessary to fully determine whether the parallel amino acids could affect functional changes, and consequently, how these changes could play a role in building superfast muscles.

## Materials and Methods

### Genome-wide screen for parallel amino acid substitutions in placental mammals

Figure S1 gives an overview of the steps. While we initially used the ortholog data from various platforms such as OrthoDB [51], OrthoMCL [52], and MetaPhOrs [53], we found that these protein sets often contain both orthologous and paralogous proteins (that could lead to spurious parallel substitutions), as well as low coverage of complete sets of one-to-one orthologs. Since high quality sets of one-to-one orthologs are essential for a genome-wide screen, we used Ensembl Compara (release 75) [16] to obtain comprehensive and accurate lists of one-to-one orthologous genes between human and other mammals and retrieved the corresponding protein sequences. We removed ambiguous amino acids (letter X) from the sequences and only kept sequences where the length is within 80-120% relative to the human or mouse sequence. All ortholog sets that contained less than 15 sequences were discarded. This resulted in 14,406 sets.

In order to reconstruct ancestral protein sequences, we first aligned each ortholog set using Prank (version 10603, parameters:+F) with the mammalian phylogeny (Figure S3) as a fixed guide tree [54], since the phylogeny-aware Prank alignment method was reported to consistently outperform other methods [55, 56]. Second, to focus on protein regions with a higher chance of being reliably aligned, we automatically removed poorly aligned regions and spurious sequences from each alignment by applying trimAl (version 1.4.rev15, “gappyout” trimming mode) [57]. This alignment trimming was shown to improve the accuracy in phylogenetic reconstructions [57].

Third, we estimated ancestral sequences using either Maximum Likelihood and a Bayesian Monte Carlo Markov Chain sampling approach. For Maximum Likelihood, ancestral reconstruction was split into two separate steps to improve computational runtime. Using RAxML (version 7.9.5) [58, 59], we first estimated for each alignment the branch lengths in a tree whose topology is constrained by the mammalian phylogeny (RAxML parameter “–g”). Then, we used Lazarus (version 2.79), which acts a wrapper for the codeML functions in PAML [60, 61], to estimate ancestral sequences using the fixed tree with estimated branch lengths as input. For both RAxML and PAML, we used the LG amino acid substitution model and its amino acid frequencies [62] and allowed for rate heterogeneity (RAxML parameter “–m PROTCATLG”, Lazarus parameter “fixed_asrv=False, model=LG”). For the Bayesian approach, we used PhyloBayes (version 4.1b) [63], constraining the tree topology by the mammalian phylogeny (Figure S3) and allowing PhyloBayes to estimate branch lengths using the default CAT substitution model [64]. Two independent Monte Carlo Markov chains were then run, sampling every 100 steps until both chains converged. Based on the PhyloBayes manual, an acceptable convergence level is reached when the maximal difference between the observed bipartitions (maxdiff) is less than 0.3 and the minimum effective sample size is larger than 50. After convergence, we proceeded to carry out ancestral reconstruction, with a burn-in of the first 100 steps and sampling every 10 steps. Since Bayesian approaches can have long runtimes (in total 390 CPU days for 14,396 proteins), we excluded proteins for which the chains did not converge after 2 weeks of runtime, which was the case for only 10 out of the 14,406 proteins. Finally, for each alignment with its reconstructed ancestral sequences that are assigned to each node in the tree, we considered all internal and terminal branches where an amino acid substitution occurred and identified all pairs of branches that exhibit exactly the same (parallel) amino acid substitution.

### Identifying radical substitutions in conserved positions

Given an observed parallel substitution between a pair of branches, we classified it as a radical amino acid substitution if the ancestral and the derived amino acid belong to a different physicochemical group (Table S2). A parallel substitution in a conserved position satisfies the following two criteria. First, all species that descend from both branches share the derived amino acid. Second, at least 90% of the species outside of the two convergent lineages share the ancestral amino acid (Figure S2). By filtering for radical substitutions in conserved positions, we enrich for parallel substitutions that are more likely to affect protein function, regardless of whether these amino acid substitutions involve a single or more than one nucleotide mutation. All primary data (alignments, parallel substitutions and their classification) are available at https://bds.mpi-cbg.de/hillerlab/ParallelEvolution/.

### Parallel substitutions in fast-and slow-twitch muscle fiber proteins

Fast-and slow-twitch muscle fiber proteins were taken from Dataset S1 in [24], which used gene expression data from isolated mouse fast-twitch and slow-twitch muscle fibers to obtain sarcomeric and Ca^2+^ signaling genes with a higher expression level in fast-twitch (slow-twitch) fibers, and from [23], which specifies the fiber type expression of myosin-associated genes. We then iterated over all pairs of independent branches, counted the number of radical parallel amino acid substitutions in conserved positions and plotted a histogram of this number. The branches in an independent pair do not share a direct common ancestor and either branch is not a descendant of the other.

### RNA-seq in *P. parnellii* muscle tissue

The use of bats complies with all current laws for animal use and experimentation in Germany, and with the Declaration of Helsinki. Anterior cricothyroid and breast muscle tissue were extracted from three adult female *Pteronotus parnellii* bats, kept in a colony in our animal house. The bats were euthanized through an overdose of Narcoren (Merial GmbH, solution contained 400 mg/kg pentobarbital) injected intraperitoneally. The anterior cricothyroid muscle was anatomically defined according to [65]. The muscle tissue was excised using surgical instruments that had been cleaned with RNAseZAP and was immediately transferred into RNAlater stabilization buffer (Qiagen).

Muscle tissue samples were approximately 1 mg and were homogenized using a pestle grinder, followed by total RNA isolation according to a standard TRIzol isolation protocol. The amount of isolated total RNA was measured with the NanoDrop (Thermo Scientific) and approximately 15 µg of RNA were found in each sample. RNA integrity was assessed using a Bioanalyzer 2100 (Agilent Technologies) and samples with RNA integrity number (RIN, a measurement of RNA degradation [66]) > 7 were selected for RNA sequencing. An Illumina HiSeq 2500 was used to generate 75 bp paired-end reads with insert size of approximately 250 bp.

### *De novo* assembly of transcriptome

Prior to transcriptome assembly, adapters and low-quality reads (minimum read length: 35 bp) were removed using Cutadapt (http://code.google.com/p/cutadapt/). Because the available *P. parnellii* genome [4] is highly fragmented, we used the Trinity (v 2.0.6) de novo assembly package [67] to assemble the *P. parnellii* transcriptome. Highly similar transcripts were collapsed using CD-Hit [68]. Annotation of the transcriptome was carried out using Blast+ against human cDNA sequences from Ensembl (version 86). We kept the best Blast hit with the lowest E-value and highest percent identity, which associates each transcript to a human Ensembl gene.

### Assembly of myosin heavy chain transcripts

To check assembly correctness, we used the UCSC genome browser’s Blat [69, 70] to align the assembled *P. parnellii* transcripts against the high-quality genome assembly of the little brown bat (myoLuc2). The longest transcripts of *Casq1, Atp2a1, Myl1* and their paralogs align with >90% identity to the orthologous gene according to the Ensembl and CESAR annotation [71, 72], showing that they are correctly assembled. In contrast, we found Myh2 to be misassembled as its transcript aligns to various parts of different myosin heavy chain genes. This misassembly is caused by the very high sequence similarity between the coding exons of the different myosin heavy chain genes, which is likely due to purifying selection.

Therefore, we used the following rationale to manually assemble the different myosin heavy chain (*Myh*) genes. The different fast-twitch fiber *Myh* genes were already present before the divergence of the amniote ancestor approximately 310 million years ago. Thus, intronic sequences that typically evolve neutrally will be diverged between *Myh2* and other paralogs. In contrast, speciation between the little brown bat and *P. parnellii* happened much more recently around 53 million years ago [73]. Consequently, the introns between the *Myh2* ortholog of both species evolved neutrally for a much shorter amount of time and thus still align (Figure S9A). Therefore, the amount of aligning intronic sequence distinguishes a genomic locus encoding an orthologous *Myh* gene from a locus that encodes a paralogous *Myh* gene.

To quantify the amount of aligning intronic sequence, we built alignment chains [74] between the little brown bat genome (myoLuc2, reference) and the *P. parnellii* genome. Then, we selected the alignment chains that best align the intronic regions to obtain *P. parnellii* loci encoding parts of an orthologous *Myh* gene (Figure S9B). By extracting the aligning exons from these loci, we manually assembled the *Myh1/2/4/6* genes. Blat of the translated transcript sequences against the little brown bat genome shows clear alignments to the orthologous genes (Ensembl and CESAR annotation), confirming the correct assembly of these four *Myh* genes. These assembled transcripts were used to quantify *Myh* expression below.

### Quantification of gene expression

We used the utility scripts provided by Trinity to automate the alignment of the reads from each tissue to the transcriptome using Bowtie2 (v 2.2.0) [75]. Subsequently, the gene expression level based on the aligned reads was estimated using RSEM (v 1.2.31) [76]. Normalization across samples was applied using the trimmed means of M-values [77]. Differential expression analysis was performed using the DESeq2 package [78].

### Functional experiments of mouse (*Mus musculus*), dolphin (*Tursiops truncates*) and microbat (*Myotis lucifugus*) calsequestrin 1

#### Protein Expression and Purification

Rosetta(DE3)pLysS E. coli cells containing the Casq1 vectors from were grown in LB media at 37 °C until an OD600 of 0.6 was reached. Next the incubation temperature was reduced to 25 °C and the cells were induced with 0.5 mM isopropyl β–D–1-thiogalactopyranoside (IPTG) and left to grow overnight. Following induction, the cells were harvested by centrifugation at 4,000g for 30 mins and then froze at −20°C for later use.

For protein purification, cell pellets were suspended in 20 mM tris, 0.5 g/L NaN3, pH 7.5 and sonicated at 8000 rpm using a 450 Sonifier^®^ (Branson Ultrasonics), until they reached apparent homogeneity. The resulting lysate was clarified by centrifugation at 20,000 × g and loaded onto a GE Healthcare AKTA^TM^ pure FPLC with an attached Toyopearl DEAE-650M (Tosoh Biosciences) column equilibrated with DEAE loading buffer (20 mM tris, 0.5 g/L NaN3, pH 7.5). Casq1 was eluted from the DEAE column between 12.5% to 25% buffer B, by running a linear gradient of Buffer A (20 mM tris, 0.5 g/L NaN3, pH 7.5) to buffer B (20 mM tris, 0.5 g/L NaN3, 2M NaCl, pH 7.5). The fractions containing Casq1 were buffer-exchanged to buffer C (5 mM sodium phosphate, 0.5 g/L NaN3, pH 6.8) and further purified with FPLC using a ceramic hydroxyapatite (HA) column (Bio-Rad Laboratories). Casq1 protein was loaded onto an HA column pre-equilibrated with buffer C. Casq1 eluted from the HA column between 50 to 100 % buffer D, by running a linear gradient of buffer C to buffer D (0.5 M sodium phosphate, 0.5 g/L NaN3, pH 6.8). For the final purification step, the HA Casq1 elution was buffer-exchanged to buffer E (20 mM tris, 0.5 g/L NaN3, pH 8.5) and loaded onto a pre-equilibrated Mono Q^TM^ column (GE Healthcare). Casq1 was eluted form the Mono Q^TM^ column by running a linear gradient from buffer E to buffer F (20 mM tris, 0.5 g/L NaN3, 2M NaCl, pH 8.5). Fractions containing Casq1 were then buffer-exchanged into Casq1 assay buffer (20 mM MOPS, 0.3 M KCl, 0.5 g/L, pH 7.2). Throughout the purification process fractions containing Casq1 were identified with SDS-PAGE. Protein concentrations were determined using the bicinchoninic acid (BCA) assay (Thermo Scientific).

#### Molecular mass determination by multiangle light scattering

We injected 100 µl of each protein at 1 mg/ml concentration, which was pre-equilibrated with chromatography running buffer (20 mM Tris-HCl pH 7.5, 300 KCl and either with or without 1 mM CaCl2), onto a Bio-Sep S-2000 column (Phenomenex). The chromatography was carried out at a flow rate of 0.5 ml/min using Acuflow series IV pump (Analytical Scientific Inst.). The eluate was passed in tandem through an UV detector (GILSON), a refractometer (Optilab DSP, Wyatt Tech.) and a multiangle laser light scattering detector (Dawn EOS, Wyatt Tech.). All the chromatography experiments were performed at 25 °C. Scattering data were analyzed using software, ASTRA (Wyatt Tech.) supplied with the instrument. Relative weight-averaged molecular masses were determined from the scattering data collected for a given condition using the Zimm fitting method, in which *K*c/R(Q)* is plotted against *sin2(Q/2)*, where Q is the scattering angle; R(Q) is the excess intensity (I) of scattered light at the angle Q; c is the concentration of the sample; and K* is a constant equal to *4π^2^n^2^(dn/dc)^2^/λ0^4^N_A_* (where n = solvent refractive index, dn/dc = refractive index increment of scattering sample, *λ_0_* = wavelength of scattered light and NA = Avogadro’s number). Extrapolation of a Zimm plot to zero angle was used to estimate the weight-averaged molecular mass.

#### Turbidity Assays

The turbidity of Casq1 solutions (i.e., absorbance at 350 nm) as a function of Ca^2+^ concentration was monitored using a Genesys 10S UV-Vis Spectrophotometer (Thermo Scientific). Assays were performed using 1.3 mL of 15 µM mouse, bat, and dolphin Casq1 in the assay buffer. Concentrated Ca^2+^ solutions (0.10 M, 0.25 M, 0.50 M, and 1.0 M) were added in 1.0 µL to 2.0 µL aliquots to the 1.3 mL Casq1 solutions in a quartz cuvette to achieve the proper calcium concentration. Upon addition of each concentrated Ca^2+^ aliquot, the samples were mixed by stirring with a small stir bar and allowed to equilibrate (dA350/dt = 0) before addition of the next aliquot. Dilutions from adding Ca^2+^ aliquots were included in data analysis.

## Acknowledgment

We are grateful to Holger Brandl for help with transcriptome analysis, Julia Jarrells and Andreas Dahl for RNA-sequencing, and Jochen Rink for helpful discussions. We thank the Computer Service Facilities of the MPI-CBG and MPI-PKS, and the Scientific Computing Facility and Gene Expression Facility of the MPI–CBG for their support. This work was supported by National Institutes of Health Grant 1R01GM11125401 and the M. J. Murdock Charitable Trust to CHK, and the Max Planck Society to MH.

